# Nanoparticles targeted to fibroblast activation protein outperform PSMA for MRI delineation of primary prostate tumours

**DOI:** 10.1101/2022.06.10.495719

**Authors:** Nicole Dmochowska, Valentina Milanova, Ramesh Mukkamala, Kwok Keung Chow, Nguyen T.H. Pham, Madduri Srinivasarao, Lisa M. Ebert, Timothy Stait-Gardner, Hien Le, Anil Shetty, Melanie Nelson, Philip S. Low, Benjamin Thierry

## Abstract

Accurate and precise delineation of gross tumour volumes remains a barrier to radiotherapy dose escalation and boost dosing in the treatment of solid tumours, such as prostate cancer. Magnetic resonance imaging of tumour molecular targets has the power to enable focal dose boosting, particularly when combined with technological advances such as MRI-LINAC. Fibroblast activation protein (FAP) is a transmembrane protein overexpressed in stromal components of >90% of epithelial carcinomas. Herein we compare targeted MRI of gold standard PSMA with FAP in the delineation of orthotopic tumours in a mouse model of prostate cancer. Control (no ligand), FAP and PSMA-targeting iron oxide nanoparticles were prepared with modification of an MRI agent (FerroTrace). Mice with orthotopic LNCaP tumours underwent T_2_-weighted 3D MRI 24 hours after intravenous injection of contrast agents. FAP and PSMA nanoparticles produced contrast enhancement on MRI when compared to control nanoparticles, which was most pronounced on the tumour periphery. FAP-targeted MRI increased the proportion of tumour contrast enhancing black pixels by 13.37% when compared to PSMA. Furthermore, analysis of changes in R2 values between healthy prostates and LNCaP tumours indicated an increase in contrast enhancing pixels in the tumour border of 15%, when targeting FAP, in contrast to PSMA This study demonstrates preclinical feasibility of PSMA and FAP-targeted MRI which can enable targeted image-guided focal therapy of localized prostate cancer.

## Background

Prostate cancer (PCa) is one of the most diagnosed malignancies in men (1) and, owing to its relatively high survival rate and frequent full gland treatment associated morbidity, is the leading cause of treatment–related years lived with disability worldwide. Along with radical prostatectomy, radiotherapy (RT) is a mainstay treatment option for all stages of disease, and is commonly delivered as a uniform dose over the entire prostate gland (2). There is mounting evidence that the risk of biochemical disease recurrence is reduced with dose escalation when greater than 80 Gy, especially in patients with intermediate or high-risk lesions. However, dose intensification of the whole gland is limited by off-target toxicities to adjacent organs at risk including the rectum, urethra, and bladder (3). Following pelvic radiotherapy, up to 30% of patients experience late grade III or IV genitourinary or gastrointestinal toxicities, with persistent long-term negative impacts on quality of life, through both mental and physical wellbeing, largely due to bowel disturbances(4, 5). Although modern RT techniques enable safer and more precise dose delivery to the prostate, treatment-associated morbidity remains a significant concern in the treatment of PCa patients.

PCa is a multifocal disease, often presenting with large primary intraprostatic lesions accompanied by smaller, well defined secondary lesions. Dominant intraprostatic lesions (index lesion) are considered to be the most aggressive, responsible for driving disease progression and prognosis, and are the most common sites of recurrence (6-8). Focal dose escalation RT approaches specifically target index lesions, balancing the risk of recurrence and patient’s quality of life. For example, the recently reported FLAME study demonstrated that focal dose escalation under the guidance of multiparametric MRI (mpMRI) could improve biochemical recurrence free survival by 7%, without impacting morbidity and quality of life (9). Early results from a phase II trial investigating MRI-guided focused ultrasound ablation for localized intermediate risk PCa also demonstrated encouraging oncologic and functional outcomes with 93% of patients disease-free at 5 months after treatment (10). In a cohort of low to intermediate risk PCa, gadolinium enhanced MRI-guided focal laser ablation showed promising early oncologic results with 17% of participants requiring additional oncologic treatment after one year of initial treatment, with no significant side effects or impact to quality of life (11).

Target delineation, a key requirement to define gross tumour volumes (GTV) in intraprostatic focal treatment approaches, is commonly referred to as the Achilles’ heel of RT. Owing to its superior soft-tissue resolution compared to computed tomography and the ability to acquire non-invasive functional imaging, mpMRI is rapidly becoming a staple tool in guiding PCa RT and more generally focal ablation. Adoption is supported by powerful technological advances such as the integration of MRI with linear accelerators (MRI-LINAC) and MRI-ultrasound (US) fusion set-ups that streamline intraoperative US image guidance with the resolution of MRI. However, mpMRI only has a moderate inter-rater agreement and is prone to a large false-positive rate and underestimation of tumour margins (12, 13). In addition, target delineation with mpMRI is not trivial, which substantially lengthens MRI-guided RT procedures and may result in greater intrafraction motion. This is further limited by magnet strength and experience of the imaging team.

A recent study demonstrated increased consensus of prostate specific membrane antigen (PSMA)-PET volumes with histology compared with mpMRI for the delineation of intraprostatic GTVs, indicating that imaging with molecular targets increases imaging specificity (14). Furthermore, GTVs derived from mpMRI significantly underestimated true tumour volumes compared to PSMA-PET. For intraprostatic radiotherapy boosting, imaging of molecular targets such as PSMA may therefore advantageously replace or be used in addition to conventional mpMRI and provides additional biological characterisation of malignant tissue for dose planning. However, PSMA-PET can intrinsically lead to GTV underestimation due to various technical aspects such as partial volume effects limiting image resolution (15, 16). In addition, PET-based target delineation suffers from the absence of integrated systems such as MRI-LINAC and MRI-US fusion. Conceptually, molecular MRI should provide improved imaging of tumoural features on a biological level over what is to date conventionally afforded by PET imaging. However, despite very extensive research in molecular MRI, especially with nanoparticulate-based contrast agents, clinical translation is yet to be demonstrated. On the other hand, most of the related research has to date focussed on diagnostic applications as opposed to the feasibility of accurately imaging and delineating primary lesions.

Besides the imaging modality itself, a key requirement for molecular MRI is the selection of a biological target specific to the tumour. PSMA is often significantly overexpressed in PCa and as such has become the gold-standard target for molecular imaging of PCa (17). Multiple PSMA ligands are now available for PET imaging with various FDA and EMA-approved PSMA-targeting probes demonstrating success in the clinic. However, loss of PSMA expression has been known to occur in neuroendocrine, androgen receptor-negative PCa, reducing the efficacy of PSMA-targeting probes in these cases. Furthermore, 5-10% of primary PCa or primary PCa lesions are PSMA negative on PET (18, 19). Therefore, imaging of PSMA is not suitable for all patients, warranting the validation of additional imaging targets in PCa (20).

Fibroblast activation protein (FAP) is an extracellular serine protease that is upregulated in stromal components of >90% of epithelial tumours but with low or undetectable expression in normal tissues. In addition, FAP expression correlates with progression and worse prognosis (21, 22). Endothelial cells can also express FAP which contributes to the regulation of tumour angiogenesis, changes in capillary morphology and microvascular reorganization (23). Specifically, FAP has been found to be expressed by tumour endothelial cells during capillary formation but is absent in the mature endothelium (24). Additionally, FAP expression correlates with a significantly increased density of microvessels (25). As such, FAP is potentially a pan-cancer marker and a promising target for molecular imaging and therapy. FAP has been shown to be upregulated in PCa, and is predominantly expressed in cancer associated fibroblasts and reactive stromal cells adjacent to carcinoma cells (26-28). Recent studies demonstrated that in a population of patients with PSMA negative tumours, intermediate to high uptake of a ^68^Ga-FAP inhibitor was observed in PCa (29, 30). Furthermore, FAP-specific PET imaging of malignancies with low PSMA expression demonstrated more precise results when compared to PSMA-PET (31). Imaging of FAP in PCa has shown promise particularly in castrate resistant prostate cancer (CRPC), with elevated FAP expression also observed in neuroendocrine prostate cancer (NEPC) and patients undergoing neoadjuvant androgen deprivation therapy (ADT) (32).

To date, the utility and significance of FAP specific MRI is yet to be assessed. To address this question, here we report on benchmarking the performance of a FAP-targeted iron oxide nanoparticulate MRI agent against PSMA in the delineation of PCa in an orthotopic mouse model. Our experimental approach builds on a previously reported FAP-ligand designed by ligating fragments of existing FAP ligands that were identified by molecular docking studies to contribute most prominently to the specificity and affinity of FAP binding. The FAP and PSMA ligands were substituted from mannose in the FerroTrace maghemite nanoparticulate MRI agent that is presently undergoing clinical trialling as a lymphotropic contrast agent. Using this novel imaging platform, we demonstrate that FAP MRI yields superior tumour contrast and coverage compared to PSMA in a preclinical model of PCa.

## Materials

The following chemicals were purchased and used as received without further purification. Trifluoroacetic acid, 1-Ethyl-3-(3-dimethylaminopropyl) carbodiimide hydrochloride (EDC.HCl), N-hydroxysuccinimide (NHS), 4-Morpholineethanesulfonic acid, 2-(N-Morpholino)ethanesulfonic acid hydrate (MES hydrate), 10X Phosphate Buffered Saline were purchased from Sigma Aldrich. Acetone and dichloromethane (DCM) we purchased from Chem-Supply. FAP binding ligand was synthesized as previously described (33). PSMA ligand Clu-CO-Lys-(tBu)_3_ ester (CAS 1025796-31-9) was synthesized by Advanced Molecular Technologies (Scoresby, Victoria, Australia).

### Analysis of FAP expression in PCa

To evaluate FAP expression in normal prostates and prostate adenocarcinoma, normalized FAP RNAseq (expected count-deseq2+1) data was obtained from the Therapeutically Applicable Research to Generate Effect Treatments initiative (TARGET)/the Cancer Genome Atlas (TCGA)/Genotype Tissue Expression (GTEx) dataset using the UCSC Xena platform (accessed 26/11/2021) (34). Corresponding patient prostate adenocarcinoma pathological T staging was obtained from the TCGA prostate adenocarcinoma (PRAD) data set (accessed 26/11/2021). Furthermore, immunohistochemistry staining of FAP in low- and high-grade prostate adenocarcinoma was obtained from the HPA059739 dataset deposited in the Human Protein Atlas (https://www.proteinatlas.org/ENSG00000078098-FAP/pathology/prostate+cancer#) (35).

### Magnetic nanoparticle synthesis

The core maghemite iron oxide (*γ*-Fe_2_O_3_) nanoparticles were synthesised via coprecipitation of iron salts in aqueous solutions and subsequent controlled oxidation using iron nitrate as previously described (36). To achieve *in vivo* stability, the nanoparticles are colloidally stabilised by block copolymers synthesised using reversible addition-fragment chain-transfer (RAFT) polymerisation. The maghemite nanoparticles and RAFT block copolymer coating have been previously developed for the preparation of a lymphotropic MRI agent (FerroTrace, Ferronova) currently undergoing clinical trialling (36). Briefly, two types of block copolymer are attached to the nanoparticle surface: a stabilising-polymer (RAFT-5-MAPC2-15AAM-3PEO) and a targeting polymer with terminal FAP or PSMA ligands (RAFT-5MAPC2-70AAM-PSMA/FAP) in a mole ratio of 70% stabilising to 30% targeting.

### Preparation of RAFT-5MAPC2-15AAM-3PEO

Polymers were prepared based on a previously published procedure with modifications (37). Methoxy triethylene glycol modified 2-{[butylsulfanyl)cabonothioyl]sulfanyl}propanoic acid (1.0 g), acrylamide (2.8 g), 4,4’-azobis(4-cyanovaleric acid) (0.050 g), dioxane (10 g) and water (10 g) were combined and dissolved before purging with nitrogen gas for 15 minutes and polymerising at 70°C for 2 hours. The mixture was allowed to cool, opened to air, then (methacryloyloxy)-ethyl]phosphonic acid (2.5 g) and 4,4’-azobis(4-cyanovaleric acid) (0.050 g) were added to the reaction mixture. The mixture was purged with nitrogen gas for 15 minutes, then heated at 70°C for 4 hours. The polymer was precipitated in acetone, repurified and collected by centrifugation.

### Preparation of RAFT-5MAPC2-70AAM-PSMA/FAP polymers

RAFT-COOH (2-(((butylthio)carbonothioyl)-thio)-propanoic acid) (0.2 g), 4,4′-azobis(4-cyanovaleric acid) (0.012 g), acrylamide (4.17 g), dioxane (7.8 g) and water (12.1 g) were combined and dissolved. The reaction was magnetically stirred at 70°C for 4 hours in an inert atmosphere. The mixture was allowed to cool, opened to air and [2-(methacryloyloxy)-ethyl]phosphonic acid (0.81 g) and 4,4’-azobis(4-cyanovaleric acid) (0.012 g) were added to the reaction mixture. The reaction was purged with nitrogen gas for 15 minutes before heating to 70°C for 4 hours under magnetic stirring. At this step the RAFT-5MAPC2-70AAM polymer was precipitated in acetone, repurified, collected by centrifugation and stored prior to conjugation with the FAP and PSMA ligands.

For preparing RAFT-5MAPC2-70AAM-PSMA, the tert-butyl ester groups of the PSMA ligand [Clu-CO-Lys-(tBu)_3_ ester] were first removed using 20% trifluoroacetic acid (TFA) in dichloromethane (DCM). Briefly, 20 mg/mL PSMA in 20% TFA in DCM was mixed for 3 hours at room temperature before vacuum drying. The residue was then dissolved in 10-20% aqueous acetic acid, transferred into a 50 mL falcon tube and a two-fold volume of chloroform was added. The reaction was mixed thoroughly before the layers were allowed to separate and the bottom organic layer containing the protective groups and non-volatile by-products was removed using a needle and syringe. The extraction of the aqueous layer was repeated twice, and it was dried under high vacuum. The residue was dissolved in glacial acetic acid, shell-freezed and lyophilized before use. The NH_2_ derivatized FAP ligand was synthesized as previously described and used for the preparation of the RAFT-5MAPC2-70AAM-FAP (31).

PSMA and FAP ligands were conjugated to the RAFT-5MAPC2-70AAM polymer using a carbodiimide coupling reaction. First, 200 mg of poly[[(methacryloyloxy)-ethyl]phosphonic acid]-*block*-poly(acrylamide), EDC.HCl (48 mg) and NHS (12 mg) were dissolved in 10 mL MES buffer (pH 5.0 – 5.5). The solution was mixed in a sonic bath for 10 minutes and the activated polymer was collected by precipitation in acetone followed by centrifugation. Second, FAP/PSMA ligands (12 mg) were dissolved in 100 μL DMSO and diluted to a total volume of 10 mL with 10X PBS buffer prior to addition to the activated polymer. The reaction mixture was stirred for 20 hours. The conjugated polymer was purified using centrifugal filters with a 3 kDa molecular weight cut off membrane. The product was diluted to a final concentration of 50 mg/mL and stored at 4°C until further use.

### Preparation of PSMA and FAP nanoparticles

17 mg of RAFT-5MAPC2-15AAM-3PEO and 20 mg of RAFT-5MAPC2-70AAM-PSMA/FAP polymers were dissolved in 2 mL of water and the pH was adjusted to 4 using 0.1M NaOH. The maghemite nanoparticles (45 mg as dried weight) were added to the polymer solution under probe sonication. After mixing for 10 minutes, the pH was adjusted to 5.0 with NaOH (0.1M). Sonication was continued for a total of 30 minutes and the pH was further adjusted stepwise to 6.0 and 7.0. The nanoparticle suspension was purified using a centrifugal filter with a 100 kDa molecular weight cut off membrane. Lastly, the nanoparticles were diluted with 3% saline to obtain a final concentration of 30 mg Fe/mL in 0.9% saline. An illustration of the PSMA/FAP nanoparticle preparation is shown in figure 2a. For the preparation of non-targeted nanoparticles the method described above was repeated except RAFT-5MAPC2-70AAM-PSMA/FAP was replaced by RAFT-5MAPC2-70AAM-COOH.

### Nanoparticle characterisation

The morphology of the coated nanoparticles was studied using transmission electron microscope (TEM; JEOL JEM-2100F-HR) equipped with a field emission gun operated at 200 kV. Images were recorded with a CCD Camera (Gatan Orius SC1000). Samples were prepared by placing a small drop of sample suspension in water onto a 200-mesh carbon-coated copper grid (ProSciTech) and subsequently evaporating the water in air.

Dynamic Light Scattering (DLS) data was recorded on a Malvern Instruments Zetasizer Nano ZS at 25°C. The hydrodynamic size of all nanoparticles suspensions was measured at a concentration of 0.1 mg Fe/mL and isotonic saline (0.9%) was used as suspending medium. Measurement duration was chosen as automatic.

Surface zeta potential data was acquired using the same instrumentation. The Smoluchowski method was used for the zeta potential measurements. The surface zeta potential of all nanoparticle suspensions was measured at a concentration of 1 mg Fe/mL and 10mM saline was used as the suspending medium.

### Cancer cell lines and culture conditions

Cells were routinely cultured in RPMI-1640 medium (LNCaP and C32) (Gibco) or Minimum Essential Medium (U87) (Gibco) supplemented with 10% fetal bovine serum (FBS) (Gibco) and 1% penicillin/streptomycin (Gibco) in a humidified atmosphere of 5% CO_2_ at 37°C. The medium was replaced every 2-3 days and cells were split once they reached 75-80% confluence using TrypLE Express (Gibco). LNCaP and U87 cells were gifted by Professor Lisa Butler and Professor Stuart Pitson, respectively.

### Flow cytometric analysis of FAP expression in U87 and C32 cells

Cells were harvested from flasks using TrypLE Select (Thermo Fisher), washed in PBS and resuspended in FACS buffer (1% bovine serum albumin (BSA) and 0.04% sodium azide in PBS). Staining for FAP was performed using either a one- or two-step process. For one-step staining, 90 μl of cells were incubated with 10 μl of anti-FAP-PE (clone 427819, R&D Systems) for 20 minutes at room temperature before washing with FACS buffer. For two-step staining, cells were incubated with mouse anti-FAP (clone 427819, R&D Systems) at a final concentration of 5 μg/ml for 20 minutes at room temperature and washed with FACS buffer, then incubated with goat anti-mouse secondary antibody conjugated to AlexaFluor488 (Thermo Fisher #A-11029) at 10 μg/ml for 20 minutes at room temperature, then washed again in FACS buffer. Samples were acquired on an LSR Fortessa or Accuri C6 flow cytometer (both BD Biosciences). Analysis was performed using FCS Express V7 Flow Research Edition (De Novo Software, Pasadena, CA, USA).

### Magnetic nanoparticle *in vitro* binding assay

To test the binding affinity of the PSMA/FAP nanoparticles, cellular uptake in cells expressing the respective targets was measured *in vitro*. LNCaP cells were seeded in a T25 cell culture flask at a density of 1 × 10^6^ cells per flask. C32 cells were seeded in a 6 well plate at a density of 3.5 × 10^5^ cells per well in the complete cell culture medium as described above. The cells were placed in a 37°C, 5% CO_2_ incubator and allowed to adhere for 24 hours. PSMA, FAP and control unconjugated nanoparticle suspensions were prepared at a concentration of 0.150 mg Fe/mL in cell culture medium. 6 mL of PSMA or control nanoparticle suspension was added to each flask containing adherent LNCaP cells or 1 mL of FAP and control nanoparticle suspension was added to each well containing adherent C32 cells. Three replicates were used to test each of the nanoparticle formulations. Cells were incubated with the nanoparticle suspensions for 24 hours in an incubator at 37°C in 5% CO_2_ humidified atmosphere. After incubation, cells were washed 3 times with PBS, detached using TrypLE Express and cell pellets were collected via centrifugation at 500 g for 5 minutes. The cell pellets were washed twice with PBS and were dried at 60°C overnight using a heating block. The dried cell pellets were digested with trace metal grade nitric and hydrochloric acid (1:1 ratio by volume). Digested samples were diluted with water to a total volume of 10 mL or 3 mL for LNCaP PSMA nanoparticles and C32 FAP nanoparticle samples, respectively. Iron concentration was measured by inductively coupled plasma mass spectroscopy (ICP-MS).

### Preclinical study in an orthotopic prostate cancer murine model

All animal experiments were approved by the Animal Ethics Committee of the South Australian Health and Medical Research Institute (SAHMRI) and were conducted in accordance with Australian National Health and Medical Research Council (NHMRC) guidelines and the Animal Welfare Act and conformed to the Australian Code for the Care and Use of Animals for Scientific Purposes. Male NOD scid gamma (NSG) aged 6-8 weeks were bred and group-housed in a specific pathogen free environment at the SAHMRI. Animals had *ad libitum* access to food and water and were clinically assessed daily. Mice were humanely euthanized via CO_2_ inhalation to remove tissues.

Intraprostatic injection of cancer cells was performed as previously described (38). Briefly, a lower midline incision was made, the bladder and prostate were externalised and 10 μL of RPMI-1640 containing 1 × 10^6^ LNCaP cells were injected into the anterior prostate using a precooled Hamilton syringe. To analyse tumour engraftment and growth, mice underwent weekly *in vivo* bioluminescence imaging (IVIS, Perkin Elmer), approximately 15-20 minutes post intraperitoneal injection with 150 mg/kg D-luciferin potassium salt.

### Magnetic Resonance Imaging

After 5-6 weeks of tumour growth, mice were intravenously injected with 40 mg Fe/kg nanoparticles in up to 100 μL of saline. Healthy mice with no tumours were used as controls. After a 24-hour period, mice were anaesthetised with 2% isoflurane in oxygen and were intraperitoneally injected with an overdose of pentobarbital prior to transcardial perfusion with 4% paraformaldehyde. Mice were postfixed in 4% paraformaldehyde prior to storage in PBS with 0.1% sodium azide. Mice underwent *ex vivo* T2-weighted multiple graded echo sequence acquired using a 16.4T Bruker Avance scanner (Bruker BioSpin, GmbH), with a 30 mm SAW coil (M2M Imaging, Brisbane, Australia) (repetition time 40 ms, echo time 10 ms, number of averages 1, slice thickness 0.1 mm) or fast low angle shot (FLASH) sequence using 9.4T Bruker BioSpec scanner (Bruker BioSpin, GmbH, repetition time 50 ms, echo time 10 ms, number of averages 1, slice thickness 0.1 mm). At the completion of scans, tumours were excised and underwent FLASH scans using a 11.7T Bruker Avance II scanner (Bruker BioSpin, GmbH, repetition time 100 ms, echo time 5.34 ms, number of averages 10, slice thickness 0.2 mm). All imaging was performed at room temperature. Horos was used to view scans (Horosproject.org, sponsored by Nimble Co LLC d/b/a Purview in Annapolis, MD USA).

Calculations of proportions of black pixels were based on protocols previously reported (39, 40). In brief, three regions of interest (ROI) were manually drawn over the prostate tumours in whole body MR scans using ImageJ software (Bethesda, Maryland, USA) (41). Pixel intensity histograms were generated for each ROI. A low intensity pixel threshold value was set, based on the visual analysis of threshold peaks.

Tumour T2 maps were generated from 16.4T and 9.4T T2-weighted MR images using an in-house code written in MATLAB (version 9.12.0 (R022a), The MathWorks, Inc., Natick, Massachusetts, United States). R2 values were calculated from T2 maps using a mono-exponential proton transverse relaxation method as previously published, using in-house code written in MATLAB(42). In agreement with previously published studies, we found negligible differences in R2 values between the two fields strengths (43, 44) and as such they have been combined in the results. T2 map tumour border ROIs were manually drawn throughout all tumour slices by an experienced MRI technician and the R2 distribution was calculated for each tumour. The resulting R2 histograms were plotted and the resulting Gaussian curves were used to calculate the change in mean R2 and differences in area under the curve.

### Haematoxylin and Eosin and Prussian Blue Staining

After *ex vivo* MR imaging, excised tumours, spleens, kidneys and livers were mounted in paraffin and 4 μm thick sections were stained with hematoxylin and eosin (H&E) or Prussian blue using standard protocols. Briefly, for Prussian blue staining sections were brought to distilled water and stained in an equal part mixture of potassium ferrocyanide and hydrochloric acid for 10 minutes, prior to washing in distilled water and counterstain with neutral red stain. For H&E staining, slides were stained using Dako automated slide stainer (Agilent, United States). Briefly, slides were brought to distilled water prior to staining with haematoxylin for 1 minute, bluing for 1 minute and eosin staining for 4.5 minutes. Images were processed using a Nanozoomer (Hamamatsu, Japan) and were visualized using ImageJ software (Bethesda, Maryland, United States). The percentage area of positive/blue Prussian Blue staining was calculated by setting colour thresholds using ImageJ software.

### Tissue Immunohistochemistry

4 μm paraffin embedded tumour serial sections were used to compare Prussian blue staining with CD31, PSMA and FAP immunohistochemistry. Staining was performed using rabbit specific HRP/DAB detection kit (Abcam) following the manufacturer’s instructions, with modifications. Briefly, after hydration through a series of ethanol, sections were treated with hydrogen peroxide blocking for 10 minutes. Antigen retrieval was performed using pH 6 10 mM citrate buffer at 98°C for 30 minutes and then incubated with 10% normal goat serum for 1 hour for protein blocking. The sections were incubated overnight at 4°C with anti-CD31 antibody (1:500, Abcam ab182981, EPR17259) or anti-FAP antibody (1:300, Abcam ab28244) or anti-PSMA antibody (1:500, Abcam ab76104, EP3253). After washing, sections were treated with biotinylated antibody and streptavidin peroxidase and developed with 3,3’-Diaminobenzidine (DAB) (all provided in kit). Images were processed using a Nanozoomer (Hamamatsu, Japan) and were visualized using ImageJ software (Bethesda, Maryland, USA).

The percentage area of positive staining was calculated by setting colour thresholds using ImageJ. Entire images have been adjusted for brightness, contrast and exposure, consistently between stains.

### Statistical analyses

Data are expressed as mean ± SEM in all cases. The significance of results were determined by unpaired t testing or one way ANOVA. Differences with a p value of less than 0.05 were considered to be statistically significant. The relationship between FAP expression and PSA levels and Gleason scores were assessed using simple linear regression. The relationship between the area of positive Prussian blue staining and CD31, FAP and PSMA staining were assessed using Spearman rank correlations using Graphpad software.

## Results

### Human FAP expression in the normal prostate and prostate adenocarcinoma

Analysis of available transcriptomic data showed a significant increase in FAP expression from normal prostate to adenocarcinoma (p<0.0001) (figure 1a). Further analysis comparing the expression between pathological T stages demonstrated significant increases from normal prostate to early stage (I and II) (p<0.0001) and to locally advanced (stage III) (p<0.0001). However, there was no additional significant increase to advanced (IV) (p>0.05) (figure 1b). FAP expression was further demonstrated by FAP immunohistochemistry in biopsies of low- and high-grade prostate adenocarcinoma (figures 1e and f). Linear correlation of Gleason score with FAP expression showed a weak positive relationship (R^2^ = 0.072) (figure 1d), whereas there was no correlation demonstrated with PSA (R^2^ = 0.005) (figure 1c).

**Figure 1.**
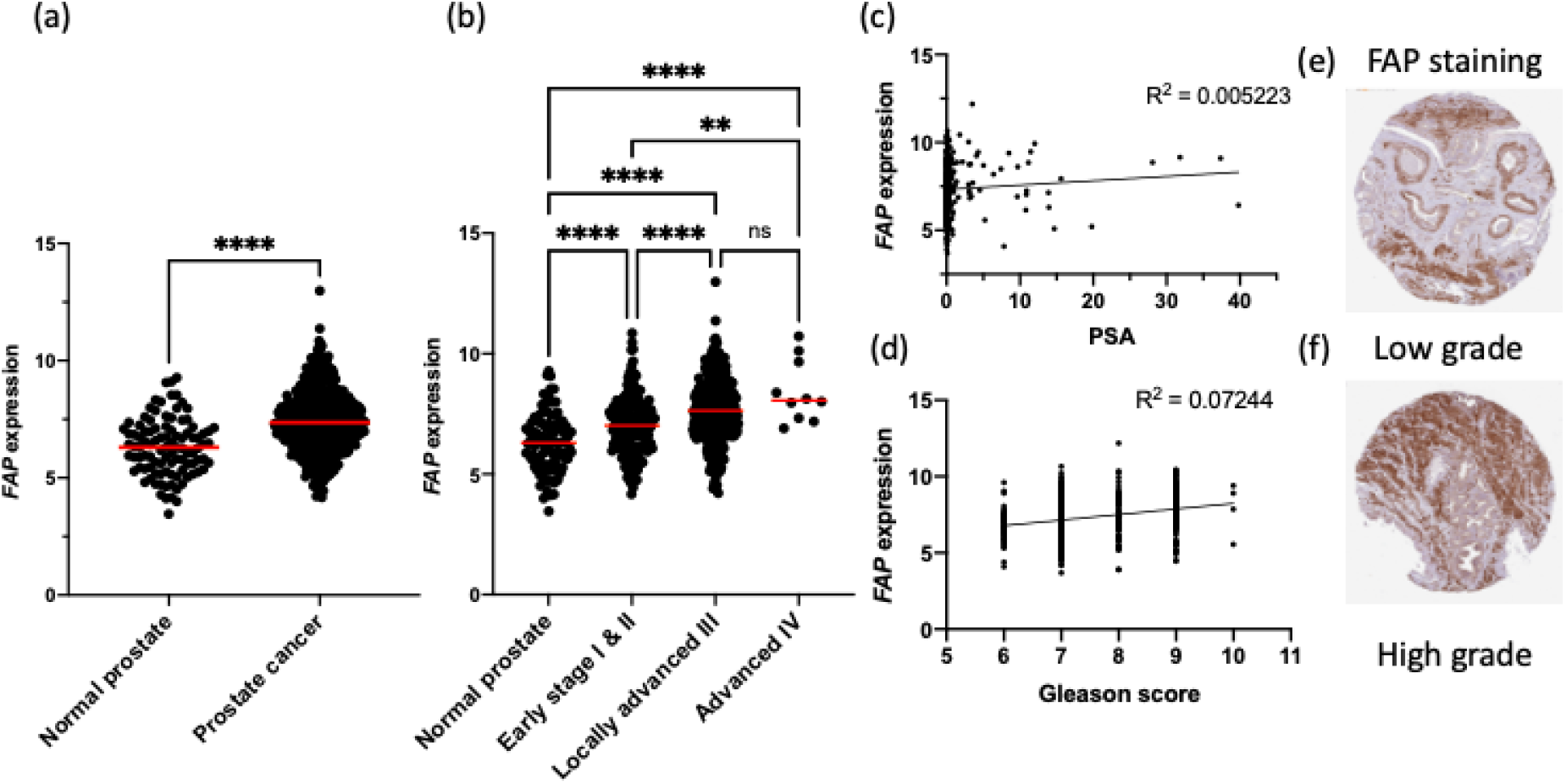
Transcriptomic analysis of *FAP* expression in clinical samples of prostate cancer. (a) *FAP* gene expression values in prostate tumours compared with normal prostate obtained from GTEX dataset. Red lines represent the medians of each group. (b) *FAP* expression in relation to pathological T staging, (c) PSA levels and (d) Gleason scores. (e) FAP immunohistochemistry in biopsies of low grade and (f) high grade prostate adenocarcinoma (images obtained from Human Protein Atlas).

### Characterization of magnetic nanoparticles

To assess the characteristics of the prepared PSMA and FAP nanoparticles, a set of physico-chemical characterisations were carried out. TEM images (figure 2b and c), taken at 60K x magnification, showed near spherical, evenly dispersed FAP and PSMA nanoparticles even when dried on a carbon film. DLS measurements of the hydrodynamic diameters confirmed that all of the coated nanoparticles have a narrow size distribution with an average hydrodynamic size of 60–65 nm when measured in physiological saline (figure 2d). All nanoparticles were also characterised with similar surface charge ranging from an average of -17 mV to -20 mV (figure 2e).

**Figure 2.**
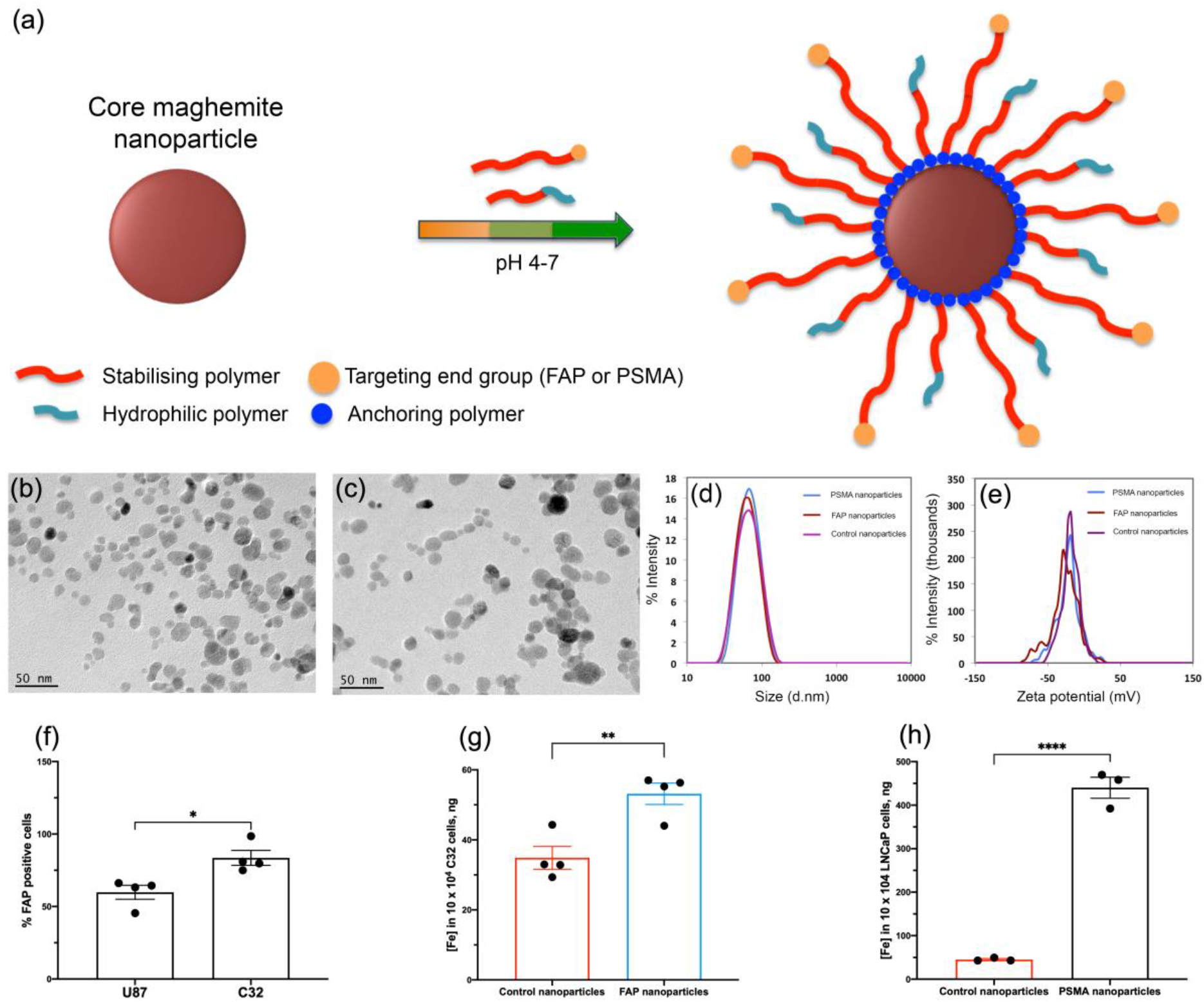
Preparation and characterization of FAP nanoparticles, PSMA nanoparticles and non-targeted control nanoparticles. (a) Schematic illustration of preparation of FAP/PSMA nanoparticles. (b, c) TEM images of FAP and PSMA nanoparticles at 60K x magnification. (d) Hydrodynamic size distribution by intensity and (e) zeta potentials of FAP and PSMA and unconjugated control nanoparticles measured by dynamic light scattering. Evaluation of *in vitro* binding activity to cells expressing molecular PSMA and FAP targets. (f) Flow cytometric analysis of proportion of FAP-positive U87 and C32 cells. (g) Cellular uptake of FAP nanoparticles and non-targeted nanoparticles in FAP-positive C32 cells. (h) Cellular uptake of PSMA nanoparticles and non-targeted nanoparticles in PSMA-positive LNCaP cells. Data presented as mean ± SEM from 3-4 replicates per condition. PSMA vs non-targeted nanoparticles, p<0.0001, t(4)= 16.306, FAP vs non-targeted nanoparticles, p< 0.0001, t(4)= 18.892, independent sample test.

### *In vitro* binding studies

The binding affinities of the prepared PSMA/FAP nanoparticles to the PSMA and FAP receptors were evaluated *in vitro* by comparing their cellular uptake in cancer cell lines which overexpress the receptors of interest. For nanoparticles used in biomedical applications, it is important to take into consideration the potential protein adsorption onto the nanoparticle’s surface which can impact the binding activity, also known as the protein corona (45, 46). For taking the formation of the protein corona into account, cellular uptake studies were performed in full cell culture media containing 10% serum and incubation time was prolonged to a 24-hour period.

First, the binding activity of PSMA nanoparticles was tested in the PSMA expressing cell line LNCaP (47). The results presented in figure 2h indicated that the non-targeted nanoparticles had a much lower cellular uptake compared to the targeted PSMA nanoparticles. Similarly, the binding activity of FAP nanoparticles was tested in the human melanoma cell line C32 as the FAP receptor is expressed in melanoma cell lines (48). Flow cytometric analysis of FAP expression demonstrated that C32 cells had approximately 25% greater proportion of FAP-positive cells in contrast to U87 cells, which had been previously demonstrated to have high levels of FAP expression (figure 2f). The targeted FAP nanoparticles had an uptake almost twice as high as the non-targeted nanoparticles (figure 2g). Considering the similar physicochemical features of the PSMA/FAP nanoparticles when compared with the unconjugated control nanoparticles, these cellular uptake data validate the chemical design and specific binding affinity of the conjugated nanoparticles to their respective targets.

### Magnetic resonance imaging of orthotopic prostate tumours

There was a significant increase in tumour contrast enhancement on T_2_-W whole body MRI after administration of both PSMA nanoparticles and FAP nanoparticles in mice bearing orthotopic LNCaP tumours compared to the unconjugated control nanoparticle group, as quantitatively demonstrated by the proportions of intratumoural black pixels (figure 3, p=0.0358 for PSMA nanoparticles and p<0.0001 for FAP nanoparticles, n=3-4 per group). FAP nanoparticle administration yielded ∼15% improvement of tumour contrast enhancement compared to PSMA nanoparticles (p=0.0004, n=4 per group). Administration of non-targeted control nanoparticles resulted in a modest, nonsignificant increase in tumour contrast relative to the no nanoparticle control group (p=0.2615, n=3 per group).

**Figure 3.**
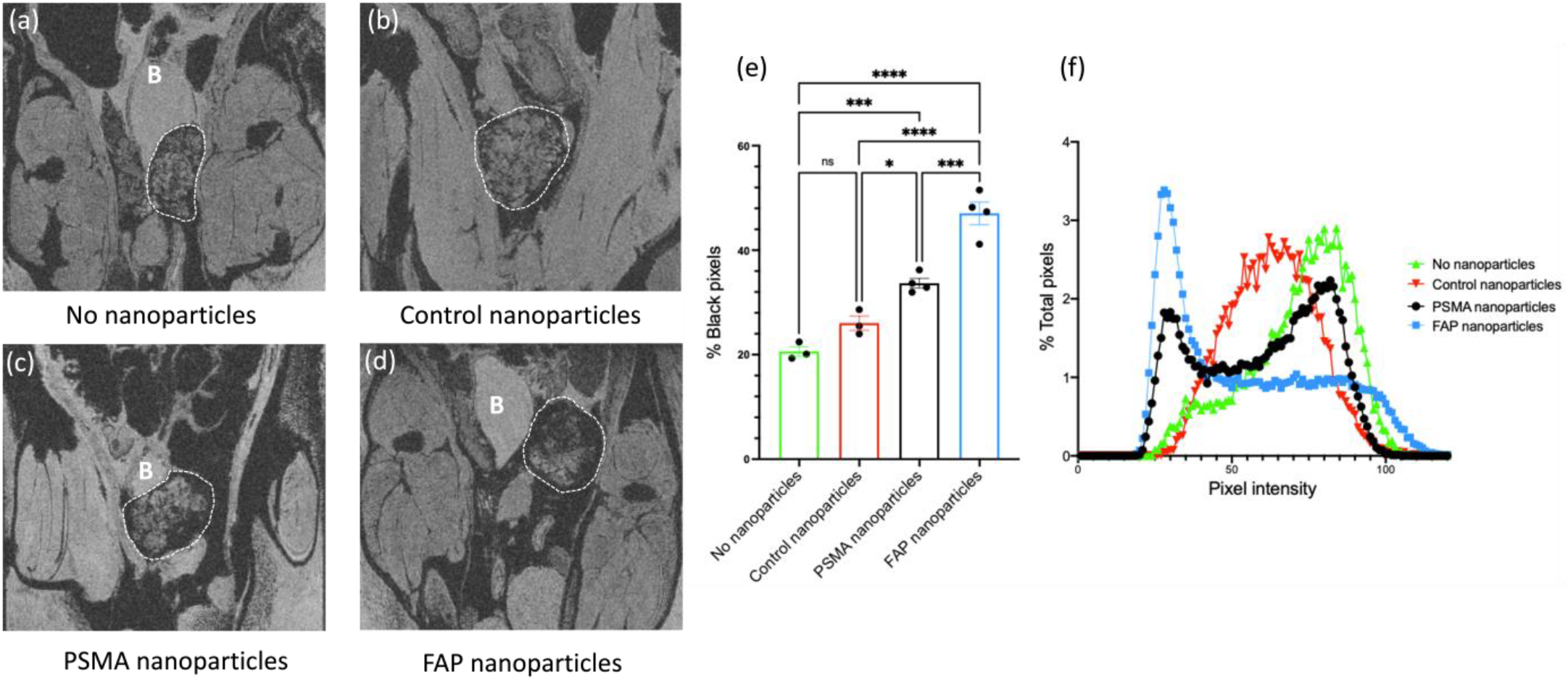
(a-d) Representative coronal view of the lower abdomen in whole body T_2_-W 16.4T MRI of mice bearing orthotopic prostate tumours 24 hours after intravenous injection with nanoparticles (40 mg Fe/kg) (representative images). PSMA nanoparticles and FAP nanoparticles enhance tumour contrast on MRI, relative to the administration of no nanoparticles and unconjugated control nanoparticles. B indicates bladder, white dotted border indicates orthotopic prostate tumour. (e) Proportion of hypointense (black) pixels on T_2_-W MRI in murine orthotopic prostate tumours 24 hours after intravenous injection of FAP and PSMA nanoparticles (data presented as mean ± SEM). (f) Representative total pixel distribution of analysed ROIs.

To better assess the potential for tumour delineation, changes in R2 distribution between prostate tumours and healthy prostates were examined. Specifically, tumour borders were compared against the healthy prostate in animals administered the same particles. Due to the overgrowth of LNCaP tumours in the prostate, these calculations were performed using separate animals. PSMA and FAP nanoparticles induced a shift in contrast in tumours (figure 4a and b), relative to control prostates, respectively. The percentage change in mean R2 relative to the healthy control with the same particles was greater in FAP nanoparticles (mean change 107%) than PSMA nanoparticles (mean change 70.14%) (figure 4c, p<0.05). More specifically, FAP nanoparticles had a greater percentage difference of area under the curve relative to the healthy control prostate in contrast to PSMA nanoparticles (figure 4d, p<0.001). Pixels in this area under the curve which is not covered by the control may contribute towards the change in contrast required for tumour delineation within the healthy prostate.

**Figure 4.**
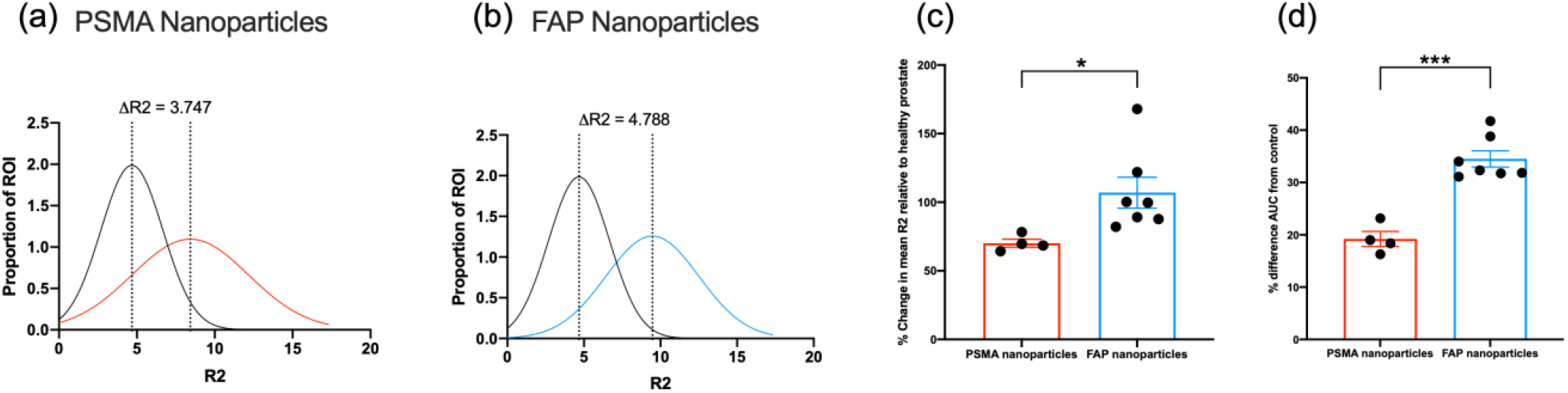
Assessment of change in R2 values in prostate tumours relative to healthy control prostates. Representative R2 traces of tumour borders after administration of PSMA nanoparticles (red line, a) and FAP nanoparticles (blue line, b) compared with healthy prostates (no tumour controls) with the same particles (black line). (c) Percentage change in mean R2 relative to healthy control, no tumour prostates with the same particles (70.14 vs 107.0, p<0.05). (d) Percentage difference of area under the curve of tumour R2 distribution curves which are not covered by the control curve (19.21% vs 34.52%, p<0.001).

### Distribution of magnetic nanoparticles in tissues

To examine the intratumoural distribution of non-heme iron after the administration of the PSMA/FAP nanoparticles, Prussian blue staining was performed on sections of LNCaP orthotopic prostate tumours and normal prostates (figure 5a). No positive staining was observed in normal prostate sections. The absence of Prussian blue staining in sections of normal prostate for mice administrated with control, FAP and PSMA nanoparticles confirmed the lack of significant nanoparticle accumulation in the absence of orthotopic tumours. LNCaP tumours also showed minimal positive basal staining. In agreement with the MRI data, modest Prussian blue staining was observed after administration of control nanoparticles. On the other hand, LNCaP tumours showed significant positive staining after the administration of PSMA and FAP nanoparticles, of which the staining was concentrated on the tumour periphery. Both PSMA and FAP nanoparticles demonstrated heterogenous intermittent intratumoural staining, some of which was suggestive of blood vessel staining, of which the frequency was greater in FAP nanoparticles compared to PSMA nanoparticles.

**Figure 5.**
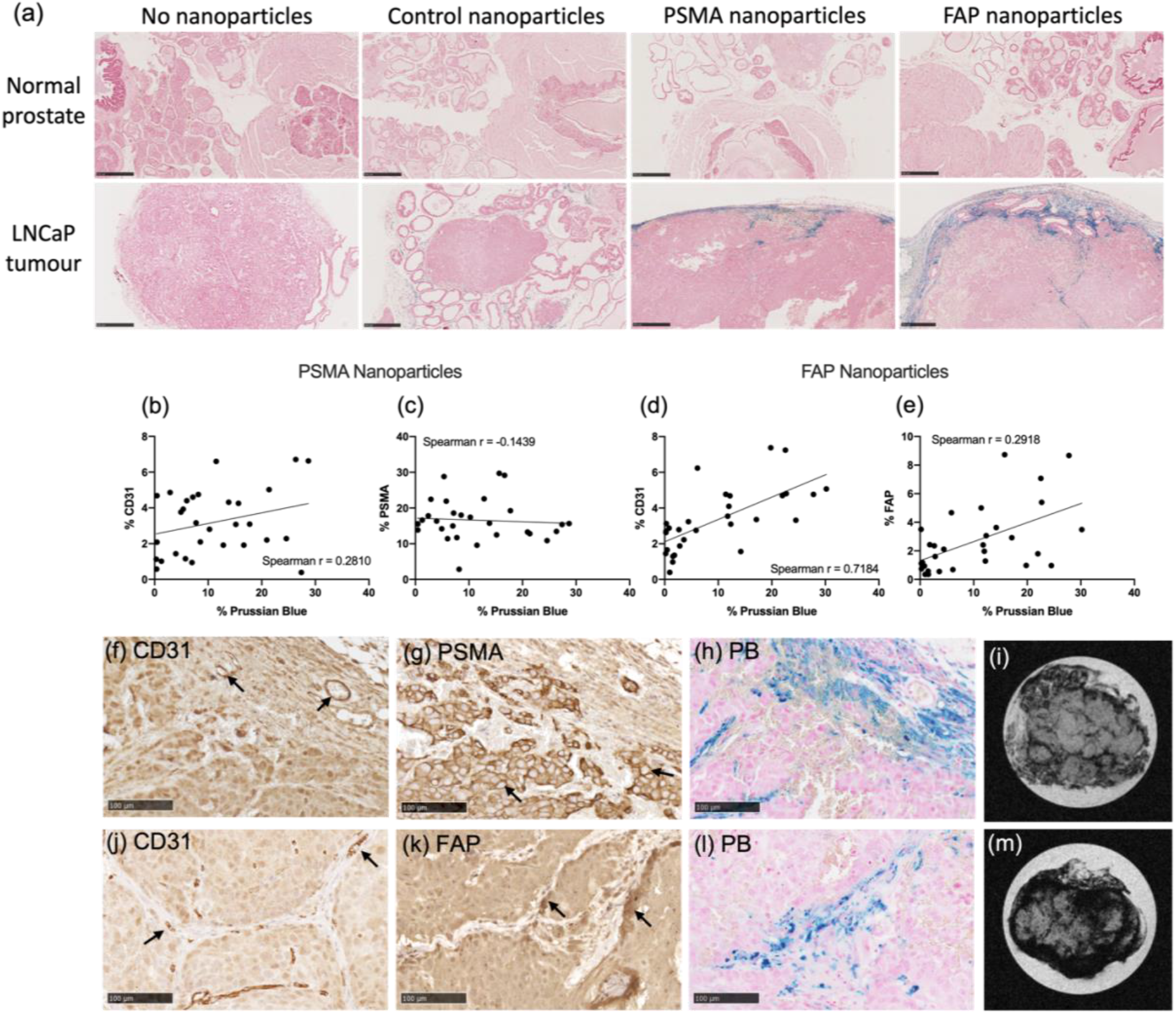
(a) Representative images of Prussian blue staining of cross sections of normal prostates and orthotopic prostate tumours after intravenous injection with no nanoparticles, control nanoparticles, PSMA nanoparticles and FAP nanoparticles (all nanoparticles 40 mg Fe/kg). All images 5x magnification, scale bar is 500 μm. (b-e) Correlation of sequential Prussian blue staining with immunohistochemistry analysis of CD31, PSMA and FAP expression in LNCaP tumours. (b and c) show correlation of colocalization of PSMA nanoparticles with expression of CD31 and PSMA, with representative images shown (f-h). (d and e) show correlation of colocalization of FAP nanoparticles with expression of CD31 and FAP, representative images shown (j-l). Representative images are 15x magnification, scale bar is 100 μm. Arrows indicate examples of positive staining. Relationship between positive Prussian blue area and PSMA or FAP or CD31 positive area is assessed by Spearman rank correlation (shown on graph). (I and m) High resolution *ex vivo* T_2_-W 11.7T MRI of murine LNCaP tumours 24 hours after intravenous injection of PSMA and FAP nanoparticles (representative images).

To better understand the distribution of the nanoparticles within tumours, positive Prussian blue stained areas were correlated with positive areas for CD31 or PSMA or FAP (figures 5b-e and representative images figures 5f-h and 5j-l). It was found that Prussian blue staining as a result of FAP nanoparticles correlated more strongly with positive area of CD31 staining compared to PSMA targeted particles (figures 5b and d). Furthermore, FAP nanoparticles had a stronger correlation with Prussian blue staining and positive staining of FAP, when compared with PSMA particles Prussian blue staining and PSMA expression (figures 5c and e). These relationships may be explained by the tumoural distribution of FAP and PSMA expression, as PSMA is expressed by LNCaP cells and therefore throughout the tumours, in contrast to FAP, which is primarily expressed by stromal components including perivascular regions. To confirm the distribution of the iron oxide nanoparticles and their effect on MRI, high resolution MR imaging of resected tumours treated with either FAP nanoparticles or PSMA nanoparticles was performed. Preferential contrast enhancement was observed at the tumour peripheries, in contrast to homogenous contrast enhancement throughout the tumours (figures 5i and m), suggesting preferential accumulation of the nanoparticles to the highly vascularised tumour periphery, in qualitative agreement with the Prussian blue staining data.

Finally, Prussian blue staining was performed to assess the distribution of iron oxide nanoparticles in the kidney, liver and spleen in orthotopic tumour mice (figure 6). Through visual assessment there was no increase in Prussian blue staining in the spleen and kidneys in all nanoparticle groups when compared to the no particle control. As expected, we observed a marginal increase in Prussian blue staining in the liver compared to the no nanoparticle control, which was consistent among all nanoparticles. Furthermore, to preliminarily assess potential acute toxicities, H&E staining was performed on the kidneys, liver and spleen of LNCaP tumour bearing mice (figure 6). No changes in tissue morphology were observed across the kidneys, liver and spleen in a control, PSMA and FAP-targeting nanoparticles, when compared to the no nanoparticle control.

**Figure 6.**
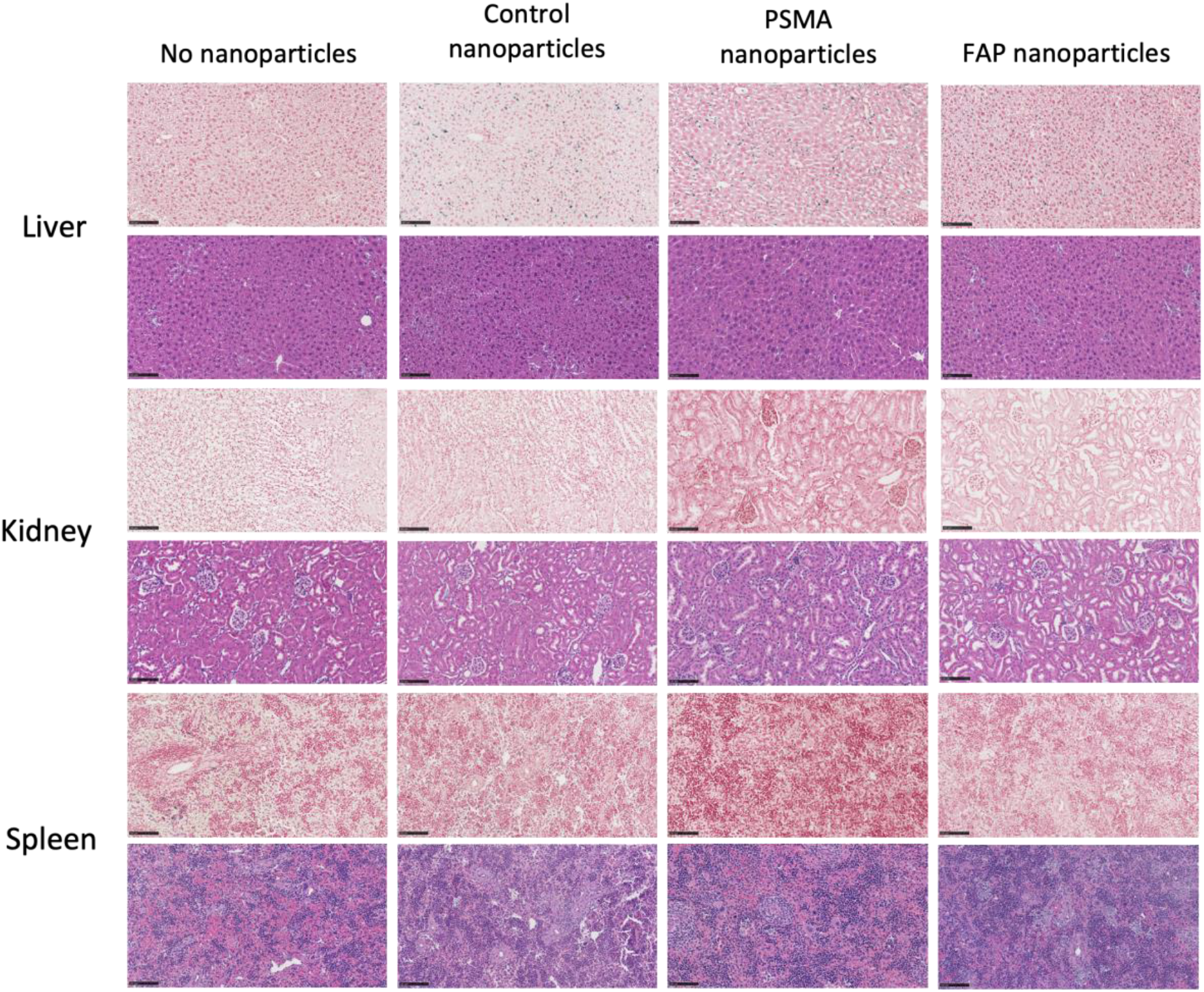
Haematoxylin and eosin (bottom row) and Prussian blue (top row) staining of liver, kidney and spleen of orthotopic LNCaP tumour mice, 24 hours after intravenous injection with nanoparticles. No changes in H&E staining were identified in the liver, kidney and spleen in all nanoparticle groups, compared to the no nanoparticle control. The kidneys and spleen did not show an increase in Prussian blue staining, relative to the no particle control, however a slight increase was observed in the liver, which was consistent across all nanoparticles. All images 20x magnification, scale bar is 100 μm.

## Discussion

Focal boost therapies of PCa aim to selectively eliminate dominant intraprostatic malignant lesions whilst sparing surrounding off-target tissues, and in doing so achieving oncological outcomes similar to those afforded by whole gland treatment but with reduced adverse effects. To enable effective focal treatment with contemporary treatment approaches including image guided RT, target volumes and safety margins must be precisely defined. Current MRI techniques such as mpMRI have yielded promising performance in providing treatment guidance but are susceptible to underestimation of the true tumour volume (49) and are time and resource demanding. This may be improved through the utilisation of molecular MRI targeting tumour markers.

Targeting of pan-cancer marker FAP has practise-redefining potential in the treatment of solid tumours such as PCa, through improved boundary delineation, possibly enabling the use of focal lesion ablative microboosts. FAP is predominantly expressed on cancer associated fibroblasts (CAFs) which reside in the tumour stroma and expression is present in virtually all solid tumours. In addition, unlike cancer cells, CAFs are not undergoing constant mutation, adding to the promising targeting qualities of FAP (33, 50, 51). Prostate tumours can have over 50% stromal component, which can be greater in cases of desmoplastic reaction (29). As such, targeting of the tumour stroma may provide increased imaging sensitivity and contrast agent retention at the intraprostatic target site compared with targeting of the tumour cells themselves. PCa metastases have also been demonstrated to be FAP positive, possibly enabling focal metastatic lesion ablative treatment, with further investigation required. In support of this train of thought, pilot studies using PSMA-PET have shown efficacy in image-guided radiotherapy of metastases for the treatment of oligorecurrent PCa (52). In another study, 1.5T MRI guided stereotactic body RT by LINAC guided by PSMA-PET produced encouraging oncological and tolerability results in the management of oligometastatic castration sensitive prostate cancer (53). Recently, FAP-targeted PET has demonstrated fair to good interobserver agreement rates in the detection of organ and lymph node metastases across a variety of tumour types (54). There are yet to be clinical studies investigating the use of molecular MRI contrast agents in image-guided RT. FAP-targeted MRI-guided radiotherapy may also prove beneficial in palliative radiotherapy, particularly in reducing pain arising from PCa bone metastases, leading to improved quality of life (55).

FAP-specific imaging may be most beneficial in late stage and clinically aggressive cancers, in which FAP expression can be more pronounced than PSMA and in cases with doubtful lesions, therapy failure and/or PSMA downregulation or negative disease (31, 56). Patients undergoing long-term ADT may have a decreased detection sensitivity on PSMA imaging due to downregulation of the receptor, although this issue may be mitigated by ceasing ADT prior to imaging studies (57). Current literature suggests that short-term ADT upregulates PSMA, suggesting that PSMA-targeted imaging is suitable in these patients.

Although various FAP and PSMA-specific PET tracers are being actively developed, PET imaging lacks the resolution that is optimal for focal radiotherapy. MR boasts superior soft tissue contrast, visualization of organ movement and the ability to monitor changes in tumour and tissue physiology, enabling improved adaptation of target volumes between RT treatment fractions (58). However, interpretation of PCa mpMRI is susceptible to significant interpretation variability as demonstrated by discordance in prostate imaging reporting and data system (PIRADS) score assignment and cancer yield by radiologists (59). MRI provides high resolution 3D morphological information which can be enhanced with the use of magnetic core nanoparticles which cause hypointense signals (darkening) on MR imaging by decreasing the relaxation times of protons in surrounding water molecules. This exogenous contrast of the tumour may lead to decreased inter-observer variability of GTV delineation. The utilisation of FAP or PSMA-targeted MRI may also improve the delineation of low-grade disease, such as Gleason 3+3 cancers which are often missed on MRI (60). Furthermore, it was found that T2-weighted MRI underestimated GTVs, longest axis and pathological tumour extent in nearly every case. Furthermore, cellular composition, density and interstitial stromal space have an effect on diffusion-weighted imaging (DWI) and derived apparent diffusion coefficient (ADC) maps, which may contribute towards the underestimation of GTVs (61).

In this study we designed, synthesized and characterized FAP and PSMA targeted iron oxide nanoparticles for molecular MRI and comparatively assessed imaging performance *in vivo* in an orthotopic model of prostate cancer. To our knowledge, this is the first report describing the use of FAP specific MRI. The ligand targeted nanoparticles are modified from a lymphotropic agent that has been validated in a large animal model and presently undergoing clinical trial (ACTRN12620000831987). This design relies on targeting and stabilising block copolymers of different lengths that enable efficient targeting using small molecule ligands as shown in the excellent binding affinity observed here *in vitro* with PSMA and FAP expressing cells.

Both FAP and PSMA targeted nanoparticles yielded improved tumour accumulation as shown by increase in MRI contrast in orthotopic tumour bearing mice compared to non-targeted nanoparticles. It is noteworthy that FAP-targeted MRI outperformed MRI targeting PSMA, the current clinical imaging benchmark, in contrast enhancement and delineation of prostate tumours in the orthotopic mouse model used here. The LNCaP tumour model was selected for this study as it commonly used for preclinical studies of PSMA specific diagnostic and therapeutic agents. Additionally, the expression of FAP in the stroma of orthotopic LNCaP tumours has been previously demonstrated as indicated by colocalization of both FAP and a-smooth muscle actin (62). The excellent imaging performance of the FAP nanoparticles observed here could be explained by the distribution of FAP expression, which predominantly occurs in the stroma and vasculature, and might therefore be more accessible to nanoparticulate imaging agents than markers expressed by tumour cells. Not surprisingly, the enhanced contrast associated to the presence of the iron oxide nanoparticles was found to be more pronounced at the periphery of the orthotopic tumours. This can be explained by the higher density of vascular structures at the periphery of such orthotopic models (63). As expected, unconjugated nanoparticles provided a modest improvement in the MRI contrast. Non-specific accumulation of nanoparticles within solid tumours is typically observed in preclinical mouse models, although the precise mechanisms are still controversial (64). Enhanced tumour contrast on MRI has also been confirmed in several clinical trials for various dextran-coated iron oxide nanoparticles (65). The ongoing NCT04682847 trial investigates the use of Ferumoxytol, a non-targeted iron oxide nanoparticle, in MRI-LINAC treatment of hepatic cancer.

Considering the pan-cancer nature of FAP expression in the stroma of solid tumours, FAP-specific MRI with rationally designed iron oxide nanoparticles may prove useful in the delineation of other solid tumours such as gastrointestinal and brain cancer. In the future, FAP specific MRI may also be useful to guide RT with proton therapies, thereby fully harnessing the uniquely precise dose deposition afforded by this technique (66).

## Acknowledgements

We acknowledge the facilities and scientific and technical assistance of the National Imaging Facility, a National Collaborative Research Infrastructure Strategy (NCRIS) capability, at SAHMRI PIRL, the University of Western Sydney and the University of Queensland.

## CRediT author statement

**Nicole Dmochowska:** conceptualization, methodology, validation, formal analysis, investigation, resources, data curation, writing – original draft, writing – review & editing, visualization, project administration. **Valentina Milanova:** conceptualization, methodology, validation, formal analysis, investigation, resources, data curation, writing – original draft, writing – review & editing. **Ramesh Mukkamala:** resources, writing – review & editing. **Kwok Keung Chow:** methodology, validation, formal analysis, data curation, writing – review & editing. **Nguyen T. H. Pham:** methodology, validation, investigation, resources, data curation. **Madduri Srinivasarao:** resources, writing – review & editing. **Lisa Ebert:** methodology, validation, formal analysis, investigation, data curation, writing – review & editing. **Timothy Stait-Gardner:** resources, data curation, writing – review & editing. **Hien Le:** writing – review & editing. **Anil Shetty:** methodology, writing – review & editing. **Melanie Nelson:** methodology, writing – review & editing. **Phil Low:** resources, writing – review & editing, funding acquisition. **Benjamin Thierry:** conceptualization, methodology, resources, writing – original draft, writing – review & editing, visualization, supervision, project administration, funding acquisition.

